# Dual-view microscopy of single-molecule dipole orientations

**DOI:** 10.1101/2025.06.17.660171

**Authors:** Yonglei Sun, Quan Wang

## Abstract

Measuring dipole orientations of fluorescent probes offers unique local structural insights of labelled biomolecules and has seen expanding applications in structural biology studies. Here we propose an alternative imaging geometry, ‘dual-view’ microscopy, for single-molecule dipole orientation measurements. We develop a protocol capable of *simultaneously* measuring absorption and emission dipole orientations of single emitters. Further, through simulation, we demonstrate that absorption dipole orientation can be accurately measured with high and uniform precision in three dimensions, significantly outperforming epifluorescence microscopy. Meanwhile the emission dipole is independently narrowed down to four possible orientations and can be uniquely determined with the co-estimated absorption dipole. Dual-view microscopy represents a new paradigm in single-molecule orientation sensing and could have unique applications in imaging under cryogenic temperatures.

**WHY IT MATTERS?:** The transition dipole orientations of fluorescent molecules bound to target biomolecules give access to the target’s local structural information, therefore can serve as unique probes in structural biology. This work proposes an alternative imaging geometry, referred to as ‘dual-view microscopy’ for more efficient three-dimensional single-molecule dipole orientation measurements compared to existing schemes. We demonstrate using simulation that our scheme outperforms conventional epifluorescence microscopy in estimation precision and can simultaneously measure absorption and emission dipole orientations independently. Dual-view microscopy represents a new paradigm that will further advance single-molecule orientation imaging as an emerging structural biology tool.

## INTRODUCTION

The interaction with electromagnetic waves of a fluorescent dye molecule can be well modeled through its electric transition dipole moment. When a dye molecule, serving as a fluorescent probe, is bound to a target biomolecule, its transition dipole orientation is dependent on the probe’s immediate structural environment. Therefore, measuring single-molecule transition dipole orientations offers complementary structural biology insights and has seen rapid growth across disciplines in recent years^1–6^. The absorption and emission dipoles of a dye molecule are not necessarily colinear as indicated by fluorescence anisotropy measurements^7^. The ability to measure *both* absorption and emission dipole orientations of single molecules is thus desirable, sometime essential in experiments.

The principles of measuring three-dimensional single-molecule absorption or emission dipole orientations are based on the fundamental physics of how a single-molecule emitter couples to electromagnetic waves. On the *absorption* part, the efficiency of excitation is determined by the alignment of absorption dipole and excitation electric field polarization. Thus, absorption dipole orientations are commonly measured by varying excitation electric field polarization either spatially or temporally^6,8,9^. On the other hand, the *emission* dipole orientation information is encoded in the dipole radiation pattern, which can be cleverly inferred by aberration or defocused imaging, phase engineering at the back focal plane, splitting emission by detection polarization, or a combination of these techniques^2,4,10–13^. Despite the diversity of single-molecule dipole orientation measurement schemes, most of them utilize the common “epi” geometry: positioning an objective lens perpendicularly to the sample for both excitation and detection (Fig. S1). We argue that the epi geometry poses fundamental limitations to single-molecule dipole orientation sensing, and propose an alternative, “dual-view” geometry. Inspired by light-sheet fluorescence microscopy^14–19^, the dual-view geometry setup positions two identical objectives orthogonally^20–22^ for exciting and collecting fluorescence emission of single molecules. We establish a measurement protocol for co-estimation of absorption and emission dipole orientations. We show by numerical simulation that our scheme achieves uniformly precise orientation measurements that outperforms the existing ones based on epi geometry.

## METHODS

### Modeling single-emitter absorption and emission with dual-view microscopy

The coordinate systems and parameters in Methods section are illustrated in Fig. S2. Let the orientation of a fixed single-molecule transition dipole positioned at the origin be represented by a unit vector *μ̂* (whether it is the absorption dipole orientation *μ̂*_*abs*_ or the emission dipole orientation *μ̂*_*em*_). Express *μ̂* in spherical coordinates under the laboratory coordinate system:

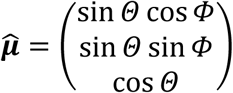

where the azimuthal angle *Φ* ∈ [−180°, 180°) and the polar angle *Θ* ∈ [0°, 180°] are the parameters of interest.

We use vectorial diffraction theory^23^ to model the efficiencies of single-molecule excitation and fluorescence collection through a tilted objective. See details in Supplemental Notes 1-3.

### Maximum likelihood estimator for absorption and emission dipole orientations

We use maximum likelihood estimation (MLE) to estimate single-molecule absorption and emission dipole orientations. Essentially, the azimuthal angle *Φ* and polar angle *Θ* of absorption/emission dipole are estimated from the recorded fluorescence photons by the proposed dual-view microscopy.

Assume a total of *n*_*tot*_ photons are partitioned into *N* portions in a particular way (as described in the Results, *N* = 6 for absorption dipole orientation estimation and *N* = 4 for emission dipole orientation estimation). We define the signal-to-background ratio *β* as the ratio of total signal photons and total background photons. Under this definition, the total number of signal photons is 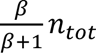 and the total number of background photons is *n*_*tot*_/(*β* + 1). Assuming the background photons are evenly distributed among all the portions, in each partitioned portion the expected number of background photons is *n*_*tot*_/[*N*(*β* + 1)].

Suppose the expected number of signal photons collected at *k*-th portion *n*_*k*,*sig*_ is partitioned by an efficiency *η*_*k*_ = *η*_*k*_(*Φ*, *Θ*), which is the excitation/fluorescence collection efficiency determined by *Φ* and *Θ* of absorption/emission dipole (Supplemental Note 3). Then

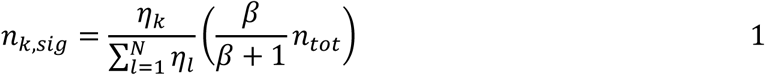

The total number of collected photons at *k*-th portion, combining both signal and background photons, is thus:

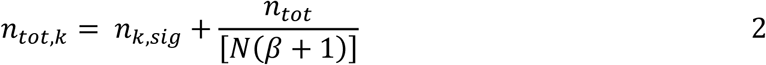

For MLE, the Poisson probability of collecting *n*_*k*_ photons given the expected mean *n*_*tot*,*k*_ at the *k*-th portion is:

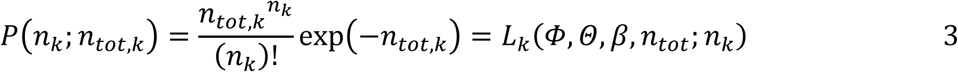

where *L*_*k*_ is the likelihood function of detecting *n*_*k*_ photons in portion *k*. Note that dependence on the single-molecule orientation (*Φ* and *Θ*) are implicitly contained in *n*_*tot*,*k*_ through Eqns. 1 and 2.

The log-likelihood of collecting *n*_1_, *n*_2_, …, *n*_*N*_ photons from the *N* portions is:

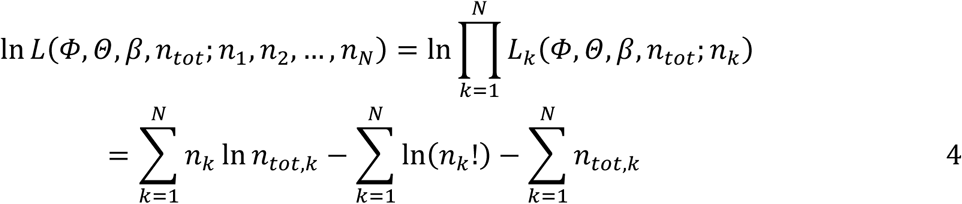

Thus, the MLE of the parameters (*Φ*, *Θ*, *β*, *n*_*tot*_) are those that maximizes ln *L*. By setting 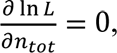 we derive the closed-form estimator 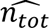 for total number of photons (note in estimation the expected total photon number *n*_*tot*_ is unknown):

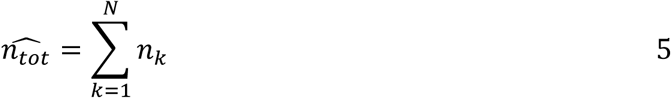

The closed-form estimators for *Φ*, *Θ*, and *β* are more difficult to obtain. We numerically search the values of *Φ*, *Θ*, and *β* that maximize ln *L*. It is equivalent to maximize the following objective function *Obj*(*Φ*, *Θ*, *β*), which is obtained by dropping terms independent of *Φ*, *Θ*, and *β* from ln *L*:

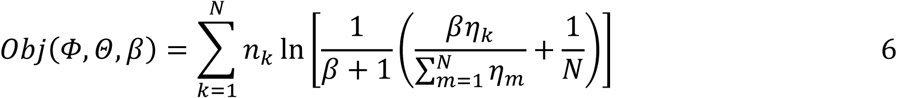

For the background-free case, i.e., *β* = ∞, the objective function *Obj*(*Φ*, *Θ*) becomes:

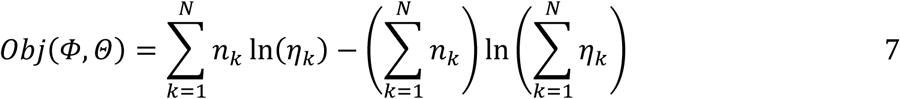

The optimization of objective function was performed using MATLAB Global Optimization Toolbox.

### Fisher information and Cramér-Rao lower bound

In order to quantify the fundamental precision bounds of the estimated azimuthal and polar angles, we calculate the Fisher information matrix from the derived likelihood function *L* (Eqn. 4). The Cramér-Rao inequality states that the inverse of Fisher information matrix (*I*(*θ*)) determines the lower bound on the variance of any unbiased estimators (*θ*), or

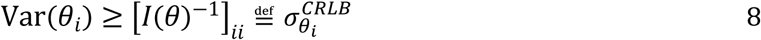

The Cramér-Rao lower bound (CRLB) 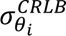 thus gives the best possible precision for estimating the parameter of interest.

Assuming the collected photons of each portion *n*_*tot*,*k*_ (*k* = 1, …, *N*) are Poisson distributed, the Fisher information matrix element corresponding to the *k*-th portion is:

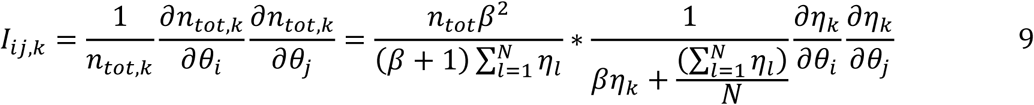

where {*θ*_*i*_, *θ*_*j*_} = {*Φ*, *Θ*}. The total Fisher information matrix element is thus:

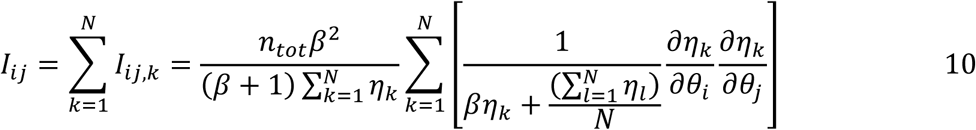

For the background-free case, i.e., *β* = ∞, the Fisher information matrix element becomes:

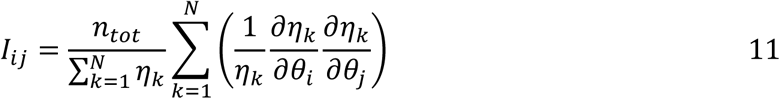

Let *I*^−1^ be the inverse matrix of Fisher information matrix. The CRLB for *Φ* and *Θ* are given by:

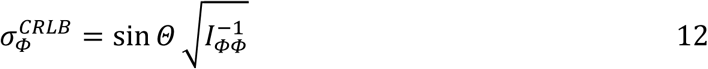

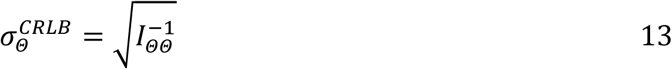

where 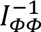 and 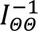 are the diagonal elements of *I*^−1^. The CRLB for *Φ* is scaled by a factor of sin *Θ* to account for the non-Euclidean nature of the rotated coordinates.^4,24^

### Generation of simulated data

In the scheme developed in the Results, we perform measurements by exciting the molecule 6 times, each time we collect the fluorescence photons from 4 avalanche photodiodes (APDs). Let *n*_*k*,*l*_ be the number of collected photons at the *k*-th (*k* = 1, 2, 3, 4, 5, 6) excitation from *l*-th (*l* = 1, 2, 3, 4) APD. To simulate *n*_*k*,*l*_, we assume they follow Poisson distribution individually, but their sum 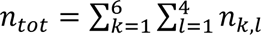 is fixed (note the degree of freedom is 24 − 1 = 23). Then the conditional distribution of *n*_*k*,*l*_ is multinomial:

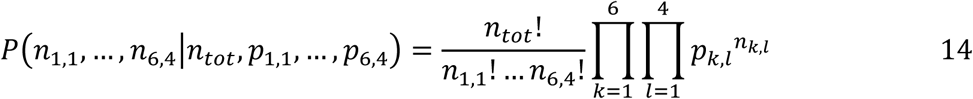

where corresponding probability *p*_*k*,*l*_ is:

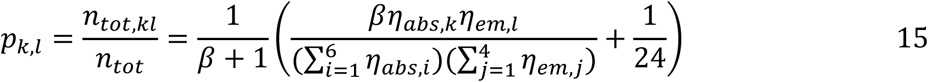

While the excitation efficiency (at the *k*-th excitation) *η*_*abs*,*k*_ and fluorescence collection efficiency (of the *l*-th APD channel) *η*_*em*,*l*_ are determined by the ground truths of absorption and emission dipole orientations respectively (Supplemental Note 3), random numbers of *n*_*k*,*l*_ are simulated from the multinomial distribution according to Eqn. 14. After simulating the 24 numbers of collected photons at each excitation from each APD, they are summed to the numbers of collected photons at each excitation *n*_*k*_ or the numbers of collected photons at each APD *n*_*l*_:

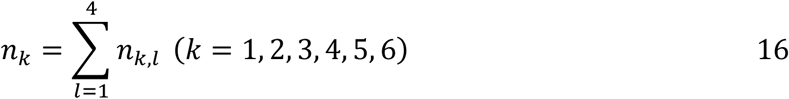

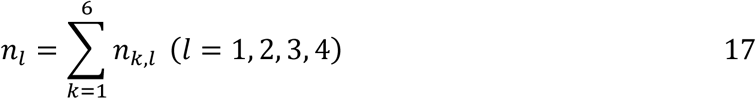

*n*_*k*_ and *n*_*l*_ are used for MLE of absorption and emission dipole orientations respectively.

## RESULTS

### Dual-view microscopy for single-molecule dipole orientation sensing

Light is a transverse wave, which means that the electric field lies in a plane perpendicular to the direction of propagation. In epi geometry (Fig. S1), this transverse plane coincides with the sample plane, so that the control of excitation polarization is limited to the sample plane. Similarly, an emission dipole perpendicular to the sample plane is not efficiently detected. We identify this geometrical constraint to fundamentally limit orientation sensing and thus propose a complementary dual-view geometry. Fig. 1a shows the proposed dual-view optical set up with two symmetric arms. In each arm an objective lens is obliquely positioned at an angle of 45° to the sample plane (i.e., the two objectives are orthogonal to each other). We assume that a single dipole emitter, of which each arm can excite and/or detect, is positioned in the sample plane and in focus for both objectives. In each arm, excitation polarization (circular insets) is controlled by a linear polarizer (LPOL). A dichroic mirror (DM) reflects the linearly polarized excitation light and transmits the fluorescence light. A polarizing beam splitter (PBS) then splits the in-plane and out-of-plane components of the fluorescence light before they are detected by separate avalanche photo diodes (APDs). Other essential components for confocal detection, such as spectral filters and confocal pinholes are omitted for simplicity. We set the *xy*-plane of the laboratory coordinate system (specified with subscript 0) coincide to the sample plane, and the *xz*-plane of the laboratory coordinate system to the plane determined by the optical axes of two objectives.

**Figure 1.**
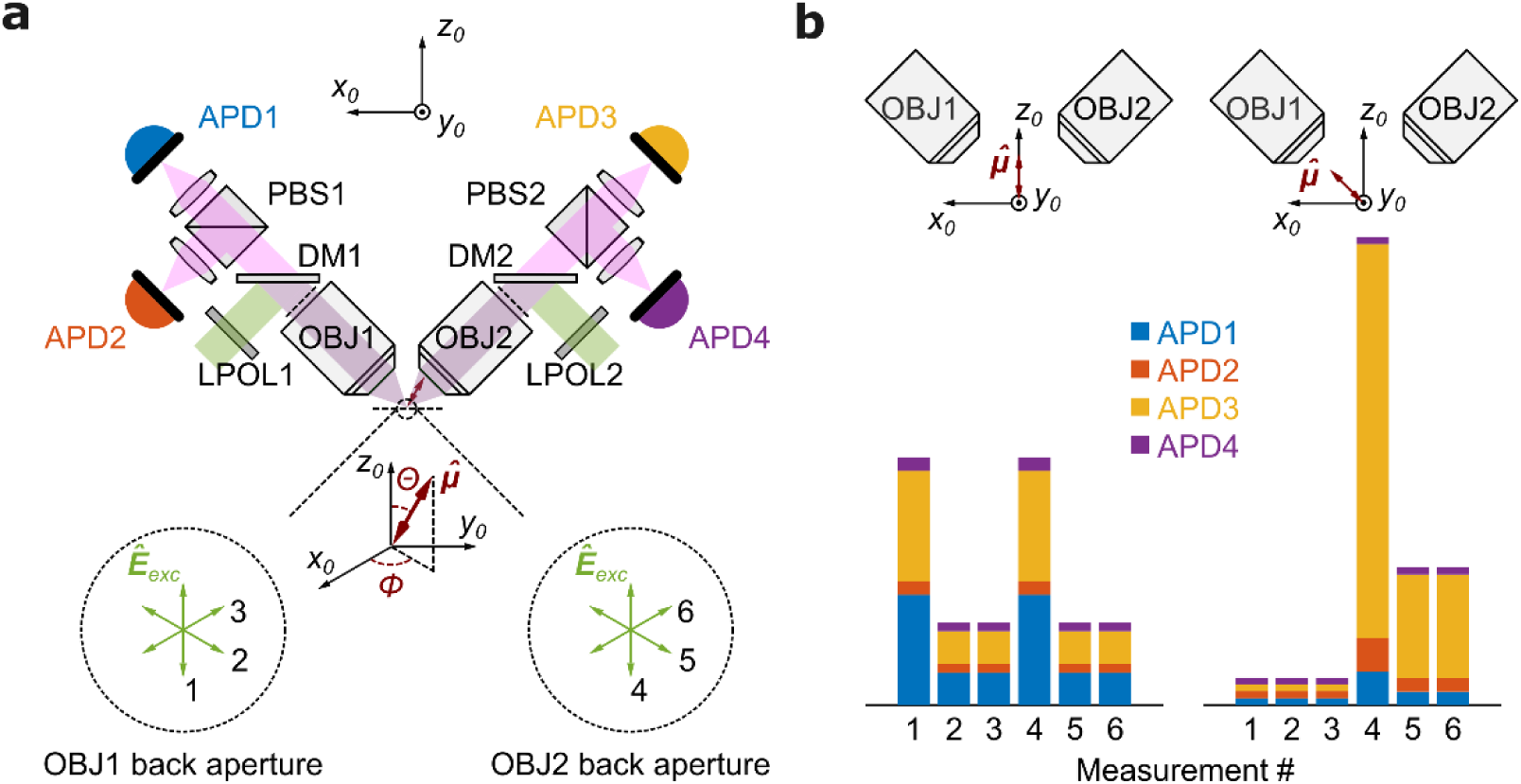
Dual-view microscopy for single-molecule dipole orientation sensing. (**a**) The proposed optical setup features two symmetric, oblique excitation/detection arms, excitation polarization modulation and polarization-resolved detection. See text for details. Insets in the two dashed circles illustrate the excitation polarization arrangements viewed from the back apertures (dashed lines before the objectives) of the respective objectives. The *xz*-plane of the laboratory coordinate system (specified with subscript 0) are determined by the optical axes of the two objectives (OBJ1 and OBJ2) from the two symmetric arms. The three-dimensional orientation of a single-molecule dipole *μ̂* is fully described by the azimuthal angle *Φ* and polar angle *Θ* under the laboratory coordinate system. (**b**) Simulated signal sets at two example dipole orientations. Here, the absorption and emission dipoles are assumed colinear and represented by *μ̂* (red arrow). Each numerically labeled vertical bar represents the total signal at the denoted excitation polarization (panel a dashed circles). The color-coded portions within each bar represent the distribution of the total signal among the 4 detectors (color coded in panel a). For clarity, photon counting uncertainties are not included.

The measurement scheme proceeds as follows. We first excite the molecule 3 times from objective 1 (OBJ1), each with a distinct excitation polarization angle *φ*_*exc*_ at the back aperture of the objective (*φ*_*exc*_ = 0°, ±60°, see Fig. 1a circular insets). Note that the excitation polarizations are in-plane at the back focal plane of the objective, and due to the oblique geometric arrangement, are *not* in the sample (*xy*-) plane after focusing. At each excitation both objectives collect the fluorescence light and all 4 APDs record the fluorescence photons. We then excite the molecule another 3 times from objective 2 (OBJ2), each with a distinct excitation polarization (*φ*_*exc*_ = 0°, ±60°), now defined at the back aperture of OBJ2 and detect the fluorescence in the same way. This protocol results in a total of 24 fluorescence intensity counts (6 excitations, each with 4 detection channels), which can be used for ratiometric estimation of single-molecule absorption and emission dipole orientations.

The *absorption* dipole orientation is estimated from the fluorescence intensity contrast following 6 polarization excitations (here, the fluorescence counts obtained by 4 APDs are summed together). At the same time, the *emission* dipole orientation is inferred from the contrast of fluorescence counts among the 4 detector (APD) channels (here, for each detector channel we sum up counts from all 6 excitations). To see how this scheme can effectively estimate both absorption and emission dipoles in three dimensions, we provide two intuitive examples shown in Fig. 1b. In the first example both the absorption and emission dipoles are parallel to the *z*-axis of laboratory coordinate system (Fig. 1b left). Due to the symmetry with respect to the two objective arms, photon counts collected in the first 3 excitations are identical to those collected in the last 3 excitations. At the same time, the measured linear dichroism from the left arm (the ratio of APD1 and APD2 counts) equals that from the right arm (the ratio of APD3 and APD4). Note that this orientation is particularly challenging for epi-detection, due to the low detection efficiency. In the second example the absorption and emission dipoles are parallel to the optical axis of OBJ1, and therefore perpendicular to the optical axis of OBJ2 (Fig. 1b right). In this case, signals collected in the first 3 measurements (excited from OBJ1) are extremely weak, highlighting the difficulty to resolve on-axis orientation with a single objective. However, due to complementarity, the same dipoles lie in the transverse plane of OBJ2 and their orientations can be resolved with high contrast in the last 3 measurements (excitations through OBJ2).

### Accurate absorption dipole orientation estimation with high and uniform precision

We first develop a maximum likelihood approach for estimating the *absorption* dipole orientation. When doing so, signals from the 4 polarization sensitive detections channels are summed up and we have 6 total signal counts resulting from the excitation sequence. The azimuthal angle *Φ* and polar angle *Θ* of the absorption dipole are estimated from the contrast among the 6 signal counts (Fig. 2a) To derive the MLE for absorption dipole orientation, we model the electric field at the focus of the excitation objective using vector diffraction theory (Methods and Supplementary Note 1) and compute the excitation efficiency given the angular separation between the electric field and absorption dipole orientation (Fig. 2a). For every possible dipole orientation and a total number of detected photons, we can then calculate the expected number of detected photons collected under each excitation. A likelihood function can subsequently be derived using Poissonian photon counting statistics (Methods).

**Figure 2.**
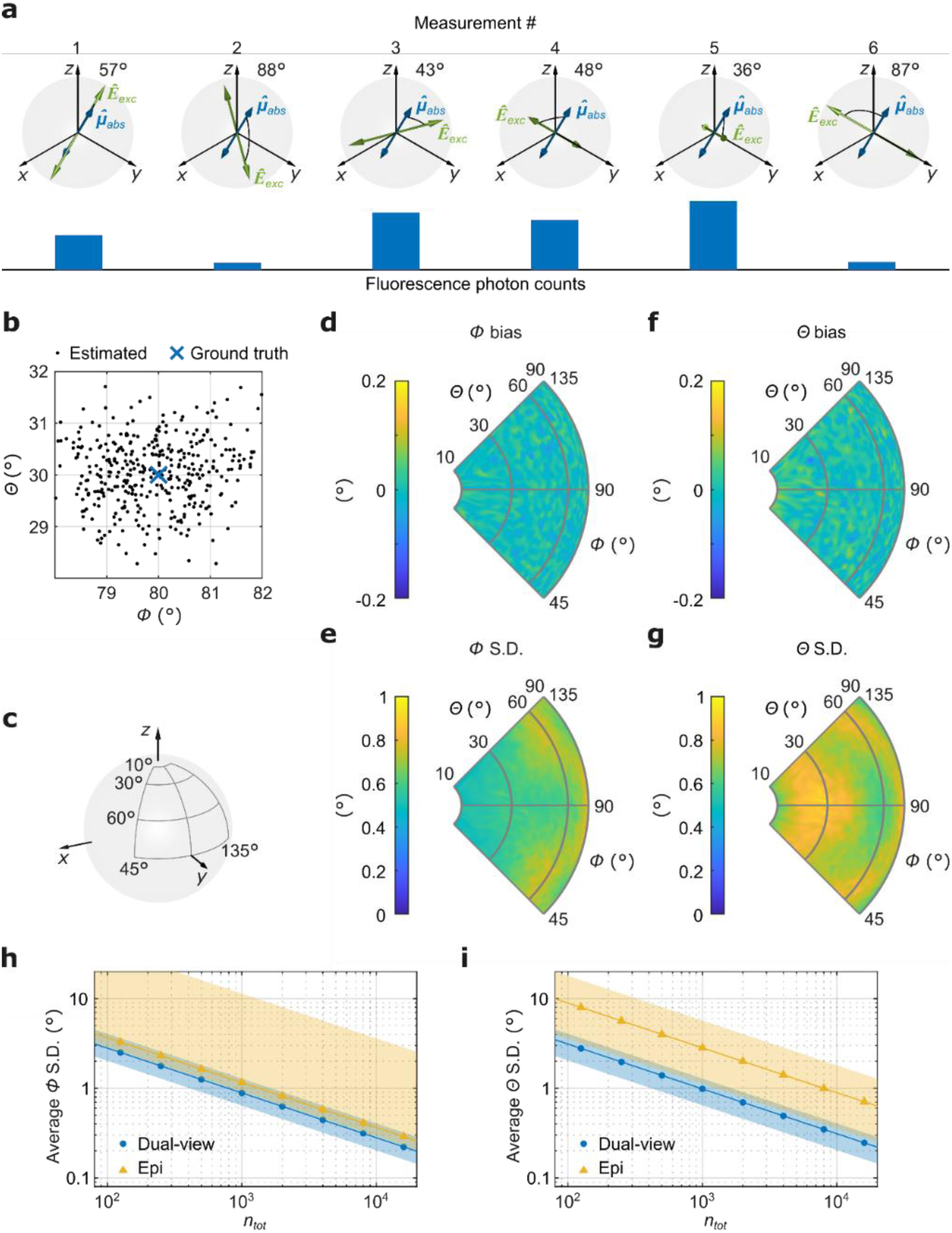
Accurate absorption dipole orientation estimation with high and uniform precision. (**a**) An example showing the angles between the absorption dipole (*Φ*_*abs*_ = 80°, *Θ*_*abs*_ = 30°) and the excitation electric field, and the corresponding expected fluorescence photon counts (blue bars). (**b**) An example of repeated estimation of absorption dipole orientations (black dots, 400 repeats) and the corresponding ground truth (blue cross) assuming *n*_*tot*_ = 5000 and *β* = 4. (**c**) Definition of the parameter space used in panels d-g. (**d-g**) Accuracy (bias) and precision (S.D.) of absorption dipole azimuthal (*Φ*) and polar (*Θ*) angle estimation in the selected parameter space shown in panel c (assuming *n*_*tot*_ = 5000 and *β* = 4). (**h, i**) Comparison of estimation precision (S.D.) using dual-view (this work, blue) and epi (yellow) microscopy schemes, as a function of total number of photons. Symbols are mean precisions over all orientations on the unit sphere. Colored shaded areas depict the spread of estimation precision over all orientations.

We first evaluate the performance on absorption dipole orientation by numerical simulations. Our focus is on estimation accuracy, precision and performance across all possible three-dimensional orientations. We assume numerical aperture *NA* = 0.5 for both objectives; the total number of collected photons *n*_*tot*_ = 5000; signal-to-background ratio *β* = 4. We also assume a separation angle *α* of 10° between absorption and emission dipole orientations, which is typical for fluorescent dye molecules.^25–27^

For a specific orientation (in this example, *Φ*_*abs*_ = 80°, *Θ*_*abs*_ = 30°), the estimated orientations, when repeated 400 times, cluster tightly around the ground truth (see Fig. 2b). The mean of repeated estimates, when compared to the ground truth, yields estimation accuracy (or bias) and the degree of scatter around the mean, quantified by standard deviations (S.D.), yields estimation precision. Regarding precision, we find that the MLE results approach the theoretical limit CRLB (Methods and Supplemental Note 4). We then evaluate the estimation accuracy and precision over the orientation space and plot a representative slice in Fig. 2d-g (approximately 1/8^th^ of the unit sphere indicated in Fig. 2c). With a realistic imaging condition (*n*_*tot*_ = 5000, *β* = 4), both the azimuthal (*Φ*) and polar (*Θ*) angles are estimated accurately (bias ≈ 0) with a precision (1 S.D.) below 1°. Most strikingly, the same levels of precision are maintained throughout the parameter space, for dipoles in the sample plane (*Θ* = 90°) as well as dipoles out of the plane (*Θ* → 0°). This is in stark contrast with epi geometry, in which out-of-plane orientations are estimated less precisely compared to in-plane orientations (see below). Furthermore, absorption dipole orientations are determined uniquely without degeneracy in most of the parameter space, except along the *Φ* = 0/180° longitude line (within *xz*-plane), where a two-fold degeneracy is present (Supplemental Note 5). Overall, these results demonstrate that our dual-view protocol achieves accurate estimation of three-dimensional absorption dipole orientations with small and uniform measurement uncertainty.

We next explore experimental determinants for estimation precision and compare to the epi microscopy^28,29^. The two critical parameters are total number of detected photons (*n*_*tot*_) and signal-to-background ratio (*β*). We calculate the precision of azimuthal and polar angles, averaged over 10,000 equally spaced orientations on the unit sphere and plot them as a function of *n*_*tot*_. The cases without background (*β* = ∞) are shown in Fig. 2h-i and the cases with *β* = 4 are shown in Fig. S6. The estimation uncertainty of both azimuthal (*Φ*) and polar (*Θ*) angles follow the 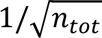 scaling. The number of photons required to achieve ∼1° uncertainty is ∼1,000 for the background-free (*β* = ∞) case and ∼2,000 for *β* = 4. As a comparison to epi-microscopy^28,29^, we plot the estimation uncertainty (CRLB, Methods and Supplemental Note 6) using the scheme introduced by Fourkas^28^. It is clearly observed from this comparison that our dual-view scheme outperforms epi-microscopy: averaged estimation precision is improved by ∼30% for the azimuthal angle and ∼3-fold for the polar angle. More strikingly, dual-view microscopy achieves uniform estimation precision (blue shaded bands) across all orientations, while epi-microscopy exhibits large precision variations (yellow shaded bands), particularly due to large uncertainties for orientations near the poles.

### Independent emission dipole orientation estimation

As described earlier, the signal contrast among the four (4) polarization-resolved detectors contains information on the *emission* dipole orientation. In a manner analogous to absorption dipole estimation, we construct the MLE for emission dipole orientation and use simulation to characterize its performance. Interestingly, a certain contrast detected on the 4 polarization resolved channels do not uniquely determine the emission dipole orientation; instead, repeated estimates are scattered around 4 “degenerate” orientations (Fig. 3a and 3b). Further investigation revealed that the 4-fold degeneracy is due to the geometry of our setup. For these four emission dipole orientations, although the *x*- and *y*-polarized intensity patterns at the back focal planes of two objectives are different, the integrated intensities detected at APDs are indistinguishable (Fig. S7). In other words, only 4 ‘integrating’ detection channels used for ratiometric estimation are not sufficient to uniquely determine the emission dipole orientation. It may seem that the ambiguity in emission dipole estimation might be a limitation of our scheme, nevertheless we next show that a simple strategy can resolve the degeneracy in most cases.

**Figure 3.**
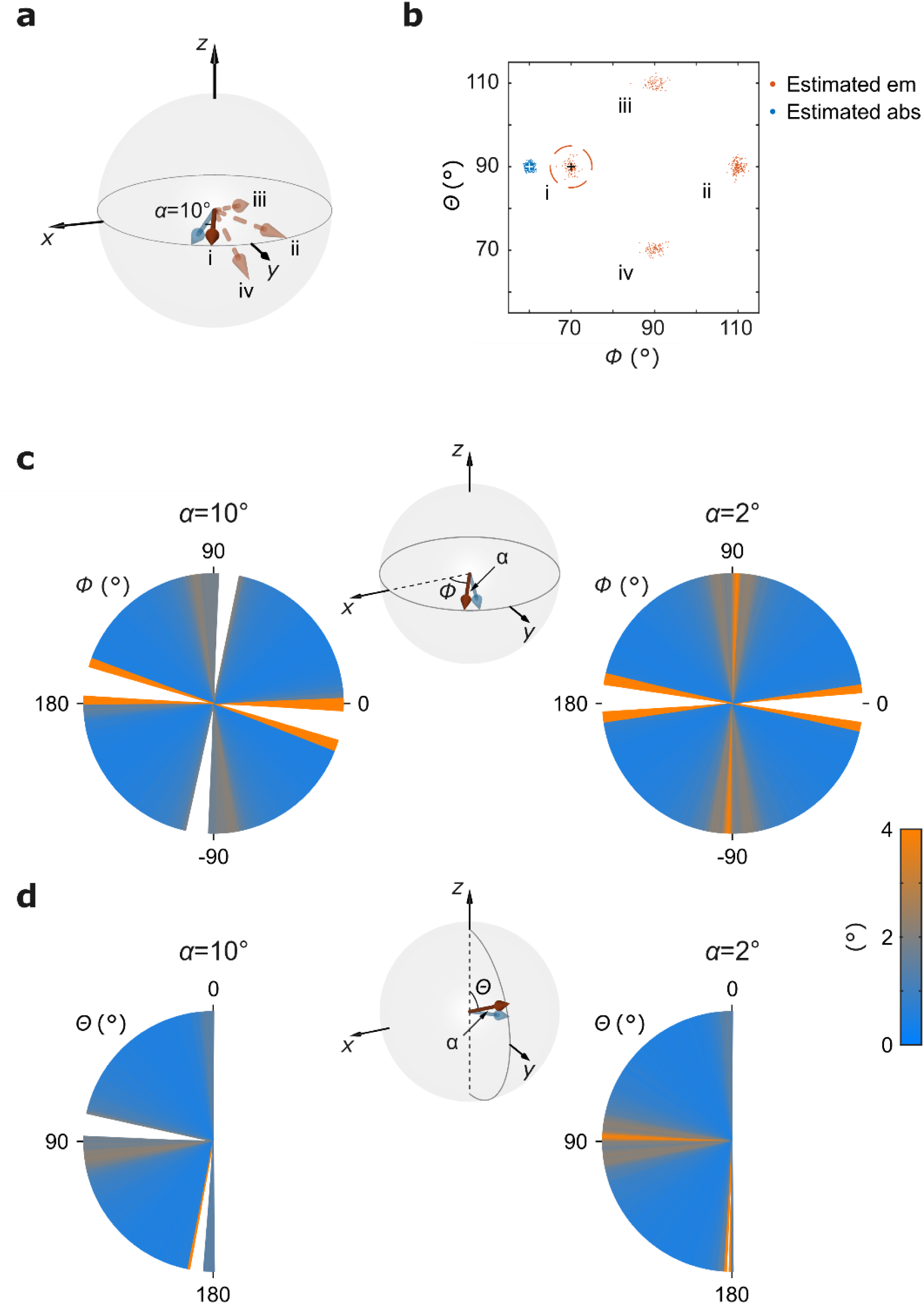
Estimation of emission dipole orientation with dual-view microscopy. (**a**) An example illustrating the 4-fold degeneracy in emission dipole estimation. Blue arrow: absorption dipole (*Φ*_*abs*_ = 60°, *Θ*_*abs*_ = 90°). Roman numerals i-iv label the 4 emission dipole orientations that are indistinguishable (degenerate) from polarization contrasts. Among those, the true emission dipole (i) is represented by a solid red arrow (*Φ*_*em*_ = 70°, *Θ*_*em*_ = 90°) the three degenerate orientations (ii, iii, and iv) are shown in semi-transparent dashed arrows. *α* is the angular separation between the absorption and emission dipoles (*α* = 10° in this example). (**b**) Resolving emission dipole degeneracy with independently measured absorption dipole orientation. With the ground truths (crosses) in panel a, the estimated emission dipole orientation (300 repeats, red dots) forms 4 clusters (Roman numerals i-iv) due to degeneracy. The estimated absorption dipole orientation is shown as blue dots (300 repeats). Degeneracy is resolved by picking the emission dipole that is closest to the absorption dipole (dashed red circle, i). (**c**) Estimation uncertainty of emission dipole orientation on the transverse (*xy*-) plane assuming *n*_*tot*_ = 5000 and *β* = 4. In this case both the absorption (blue arrow) and emission (red arrow) dipoles lie in the *xy*-plane with an angular separation of *α* between the two (middle schematic) and dipole estimation reduces to only the azimuthal angle (*Φ*). Uncertainty (of *Φ*) is shown as a color map around the equator (*Φ* from −180° to 180°). Left: *α* = 10°, right: *α* = 2°. *Φ* values with a hit rate below 90% are not shown (see text). (**d**) Estimation uncertainty of emission dipole orientation on the longitudinal (*yz*-) plane assuming *n*_*tot*_ = 5000 and *β* = 4. In this case both the absorption (blue arrow) and emission (red arrow) dipoles lie in the *yz*-plane with an angular separation of *α* between the two (middle schematic) and dipole estimation reduces to only the polar angle (*Θ*). Uncertainty (of *Θ*) is shown as a color map around the meridian plane (*Θ* from 0° to 180°). Left: *α* = 10°, right: *α* = 2°. Θ values with a hit rate below 90% are not shown (see text).

### Pick emission dipole orientation with absorption dipole orientation information

We propose a simple strategy to pick the emission dipole orientation from the four possible candidates: we perform an independent estimation of the absorption dipole orientation on the same setup as described earlier and pick the emission dipole orientation that is closest in orientation space. Fig. 3b illustrates an example: for a single molecule with non-parallel absorption and emission dipoles, we can estimate the two independently. While the absorption dipole orientation can be uniquely determined (blue), 4 possible emission dipole orientations are found (red). In this case, we pick orientation ‘i’ (red dashed circle) due to its closest proximity to the absorption dipole orientation, i.e., that it spans the smallest angle with the absorption dipole orientation. We anticipate this strategy to work well when the angle between the absorption and emission dipoles is small, which was found to be true (typical *α* ∼0-13°)^30–33^ or can be evaluated using fluorescence anisotropy experiments^7^. In general, one orientation measurement returns a single value of the emission dipole estimation. We can calculate the other three degenerate orientations based on geometry and pick the one that is closest to the estimated absorption dipole.

We next evaluate the performance of emission dipole estimation with this strategy to resolve degeneracy. For clarity and without loss of generality, we limit the evaluation to dipoles on either the transverse (*xy*-) or longitudinal (*yz*-) planes. In all cases, the absorption and emission dipoles are not parallel and differ by an angle of *α*. For a given set of orientations, we simulate the full measurement scheme (6 excitations, 24 total photon counts, Fig. 1) and estimate the absorption and emission dipole orientations independently and simultaneously. Repeated simulations on the same ground truth orientations generally result in one cluster of absorption orientations and four clusters of (degenerate) emission orientations (Fig. 3b). Degeneracy is resolved by picking the orientation closest to the estimated absorption dipole and the precision of emission dipole estimation can be evaluated.

First, we evaluate the probability that our simple degeneracy-resolving scheme returns the correct emission dipole orientation (i.e., “hit” rate, Supplemental Note 8). We find the “hit” rate is unity in most regions in the case studies. Nevertheless, in certain regions (e.g., *Φ* ∼ ± 90° and *Θ* = 90°), the hit rate drops below 90% (due to geometrical properties of the degenerate points, Supplemental Note 7). Intuitively, the sizes of these regions shrink when the angular difference (*α*) between the absorption and emission dipoles decreases.

Next, in the parameter regions where the “hit” rate is high (>90%), we benchmark the precision of emission dipole estimation in Fig. 3c and 3d. Here, the precision is quantified by the standard deviation (S.D.) of 300 repeated simulations with the degeneracy-resolving scheme, each assuming *n*_*tot*_ = 5000 and *β* = 4. Even for a relatively large angular separation (*α* = 10°, left), our scheme achieves uniform and high precision (S.D. ∼1° for 5000 total photons) in most regions of the parameter space. Only near (*Φ*∼ − 98°, −10°, 83°, 170° in panel 3c and *Θ*∼83°, 173° in panel 3d), the precision becomes worse due to reduced ability to resolve emission dipole degeneracy. These results demonstrate that our dual-view microscopy scheme can precisely measure emission dipole orientations for typical single dipole emitters.

## DISCUSSION

In this work we develop a dual-view microscopy scheme to measure single-molecule absorption and emission dipole orientations. Our scheme features two objectives arranged in a tilted and complementary geometry. The measurement protocol consists of a 6-polarization excitation sequence and polarization-resolved fluorescence photon counting. We show that signal contrasts among the 6 polarization excitations yield absorption dipole orientation via a maximum likelihood estimator. The estimates are accurate with high precision uniform across all orientations. At the same time, by sorting the same batch of detected fluorescence photons into the 4 polarization-resolved detection channels, the contrast can inform emission dipole orientation with high precision, albeit with a 4-fold degeneracy. We further show that the degeneracy can be resolved by simply picking the estimated emission dipole that has the smallest angular separation with the estimated absorption dipole. Altogether, we establish dual-view microscopy as a powerful new modality for single-molecule dipole orientation measurement with unique advantages.

Many clever schemes to measure single-molecule dipole orientations have been developed. Most of these methods are based on the epi geometry and use high *NA* objectives, together with phase processing (for example, by defocusing or patterned phase masks^12,13^) to enhance orientation sensing along the optical axis (i.e., perpendicular to the sample plane). However, due to the intrinsic limitations of epi geometry discussed earlier, these schemes struggle to achieve uniform precision in three dimensions and often result in in-plane dipoles to be measured more precisely than out-of-plane ones^20,34,35^. Our scheme is fundamentally different in that it uses geometry and two complementary perspectives to achieve three-dimensional orientation sensing with near uniform precision. No phase masks, defocusing or image processing are required, although incorporation of these elements could further enhance sensing capacity. Recently, Chandler *et al.* developed a pol-diSPIM microscope^22^, which shares many advantages to our scheme developed here. But their focus is on deconvolving the orientation information from ensembles of molecules, not on single molecules. Our measurement schemes requires ∼10^3^ signal photons to achieve 1° uncertainty of azimuthal and polar angles estimation for both absorption and emission dipole orientations, a performance comparable to existing three-dimensional state-of-the-art absorption^36,37^ or emission^4,12,34,35,38^ dipole orientation measurement schemes using immersion objectives with larger effective *NA* (typically > 1.2) values.

Another unique feature of our scheme is that absorption and emission dipoles can be estimated simultaneously. In fact, both measurements use the same batch of fluorescence photons, just grouped differently, highlighting that single fluorescence photons contain information of *both* absorption and emission dipole orientations. This unique capacity of exploiting precious single-molecule emission photons is particularly useful for measurements when both dipole orientations of molecules are required, a tall task which has only been achieved in very limited cases^37^.

We note that the current scheme also has a few limitations. First, the emission dipole orientation is not uniquely determined but has a 4-fold degeneracy. Replacing the APDs with slightly defocused array detectors (e.g., cameras or quadrant photodiode) could readily resolve this degeneracy. Second, the use of air objectives could limit the achievable signal-to-background ratio for single-molecule imaging. Nevertheless, we envision an ideal application to be imaging at cryogenic temperatures^39^, under which immersion objectives are difficult to integrate.

## ACKNOWLEDGMENTS

We thank Bok-Eum Choi, Lukas Whaley-Mayda, Jin Qian, Hoi Sung Chung and his lab for helpful discussions.

This work utilized the computational resources of the NIH HPC Biowulf cluster (https://hpc.nih.gov/). This work was supported by the Intramural Research Program of the National Institute of Diabetes and Digestive and Kidney Diseases, National Institutes of Health.

## AUTHOR CONTRIBUTIONS

Y.S. and Q.W. designed the research. Y.S. derived, implemented the model, and analyzed the data under Q.W.’s supervision. Y.S. and Q.W. wrote the manuscript.

## Supplementary Information

**Figure S1.**
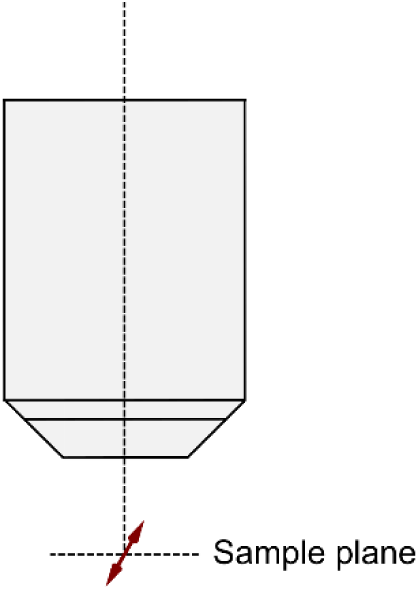
“Epi” geometry. The objective lens is positioned so that its optical axis is perpendicular to the sample plane for both excitation and detection.

**Figure S2.**
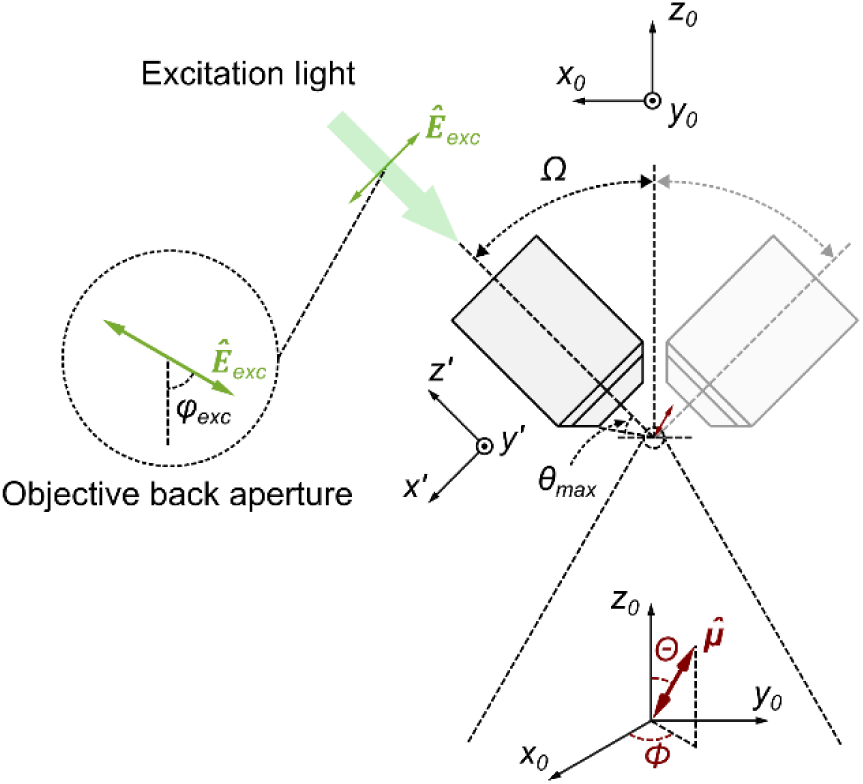
Coordinate systems and parameters for dipole orientation measurements. The laboratory coordinate system is specified with subscript 0. The rotated coordinate system specified with the prime sign comes from a right-handed rotation of an angle *Ω* about the *y*-axis of laboratory coordinate system. This rotation maps the *z*-axis of laboratory coordinate system to the optical axis of the objective. The objective is characterized by its numerical aperture *NA* = *n* sin *θ*_*max*_, where *n* (= 1) is the refractive index and *θ*_*max*_ is the half-aperture angle of the objective. The excitation polarization *Ê*_*exc*_ is fully described by the azimuthal angle *φ*_*exc*_(seen from objective back aperture) under the rotated coordinate system. The dipole orientation represented by unit vector *μ̂* is described by the azimuthal angle *Φ* and polar angle *Θ* under the laboratory coordinate system.

**Figure S3.**
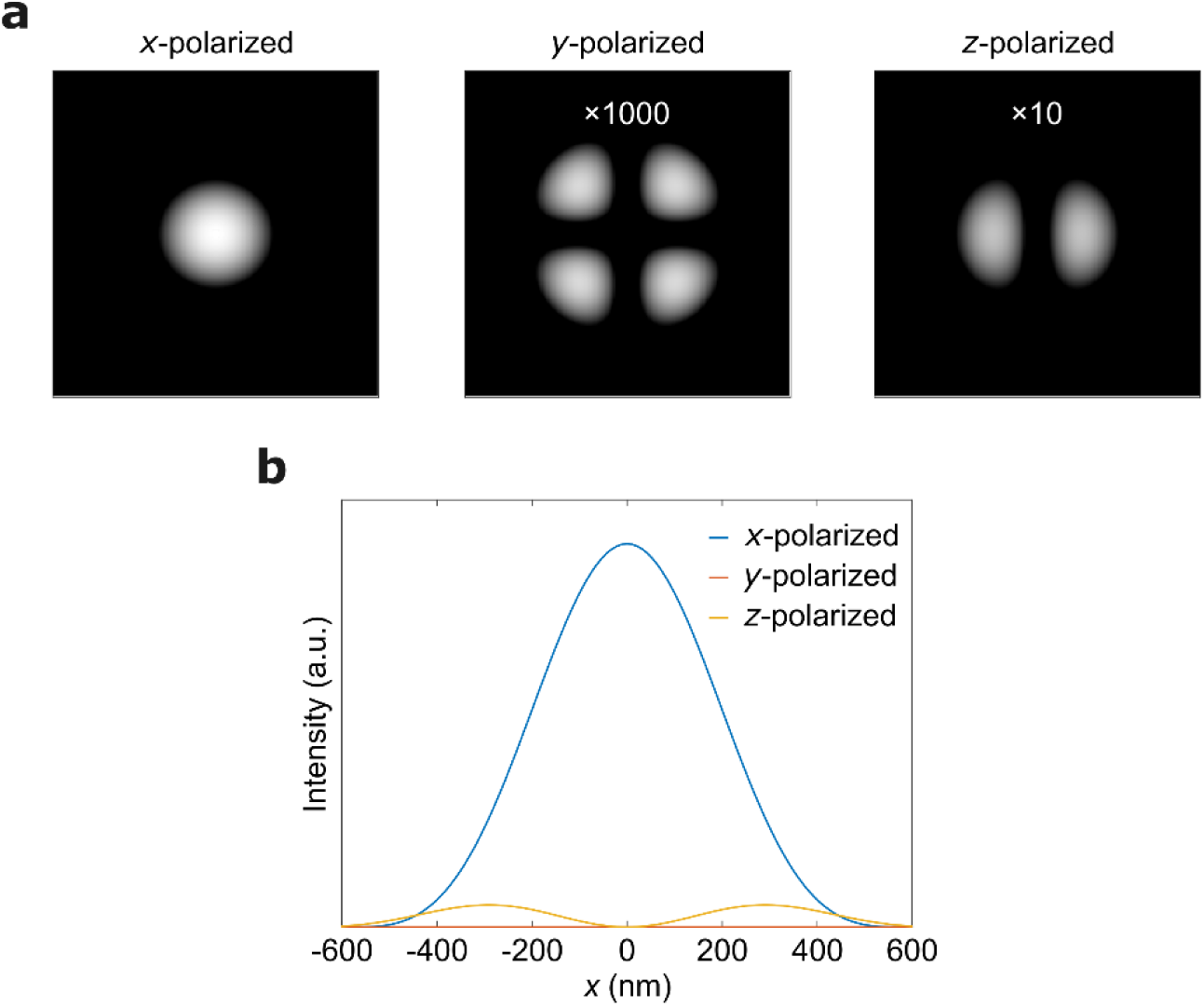
Electric field intensity distributions at the focal plane assuming the incident laser light has a wavelength of *λ* = 532 nm and is polarized along the *x*-axis. The calculation is based on vectorial diffraction theory. See details in Supplemental Note 1. (a) *x*-, *y*- and *z*-polarized electric field intensity distributions at the focal plane (region size 4*λ* × 4*λ*) assuming *NA* = 0.7. *y*- and *z*- polarized electric field intensities are amplified by 1000 and 10 times respectively. (b) *x*-, *y*- and *z*-polarized electric field intensity profiles along the *x*-axis at the focal plane.

**Figure S4.**
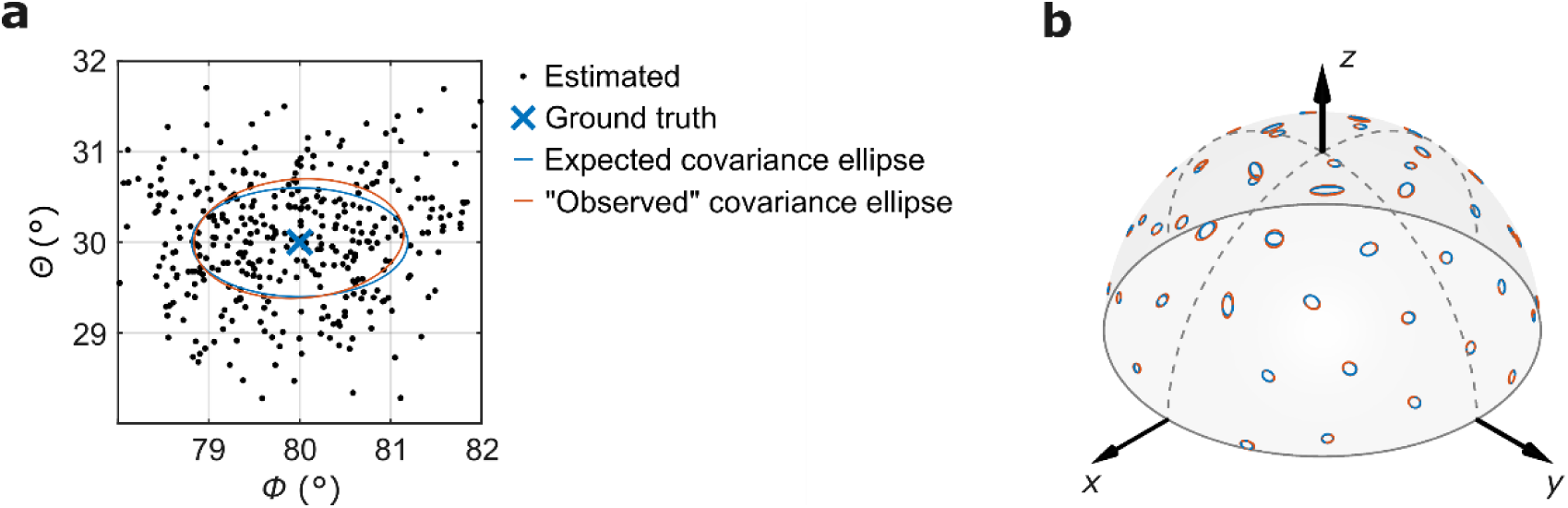
Expected and “observed” covariance ellipses comparison. (a) Expected covariance ellipse (blue) and “observed” covariance ellipse (red) calculated from the example in Fig. 2 a-b in the main text. (b) Calculated expected covariance ellipses (blue) and “observed” covariance ellipses (red) of 50 equally distanced orientations on the hemisphere assuming *n*_*tot*_ = 5000, *β* = 4. For each absorption dipole orientation, the simulation and estimation process is repeated 400 times to calculate corresponding “observed” covariance ellipse.

**Figure S5.**
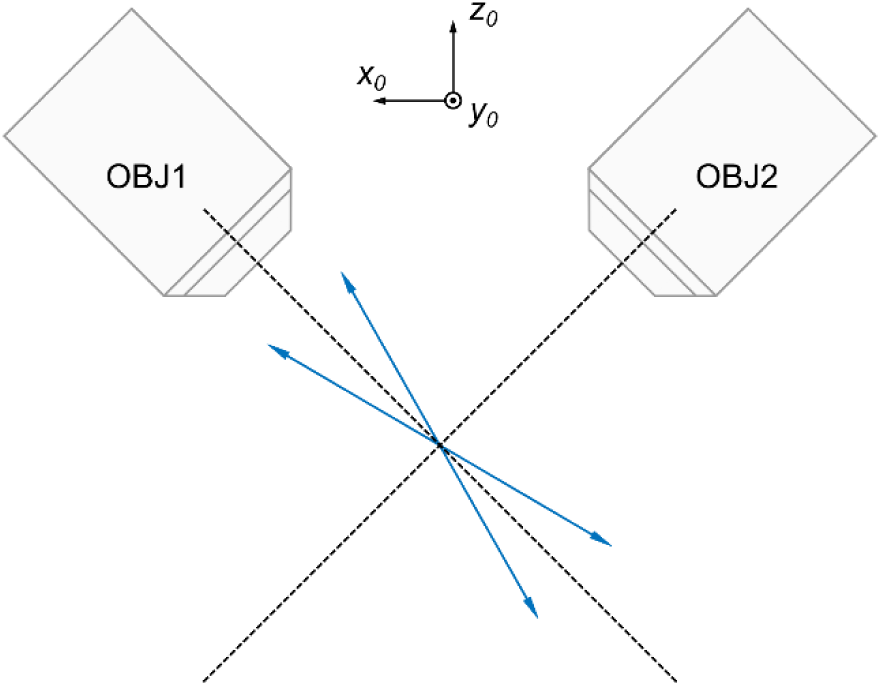
2-fold degeneracy of absorption dipole orientation estimation along the *Φ* = 0°/180° longitude line (within *xz*-plane).

**Figure S6.**
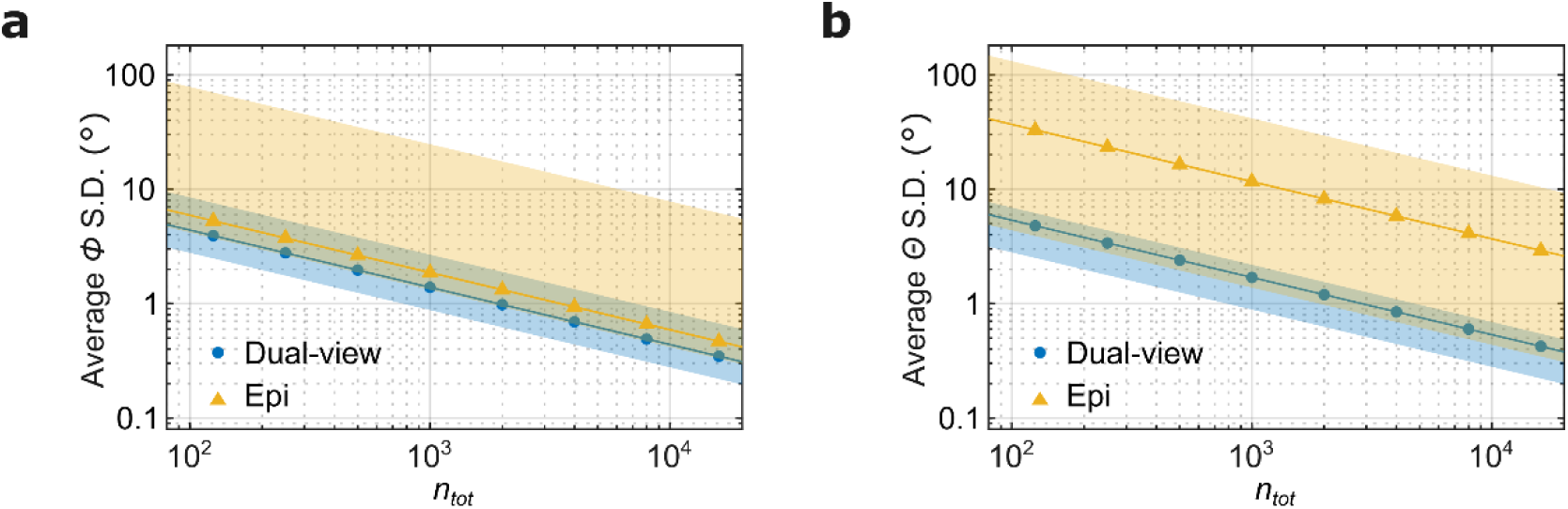
Comparison of estimation precision (S.D.) using dual-view (this work, blue) and epi (yellow) microscopy schemes, as a function of total number of photons at signal-to-background ratio *β* = 4. Symbols are mean precisions over all orientations on the unit sphere. Colored shaded areas depict the spread of estimation precision over all orientations.

**Figure S7.**
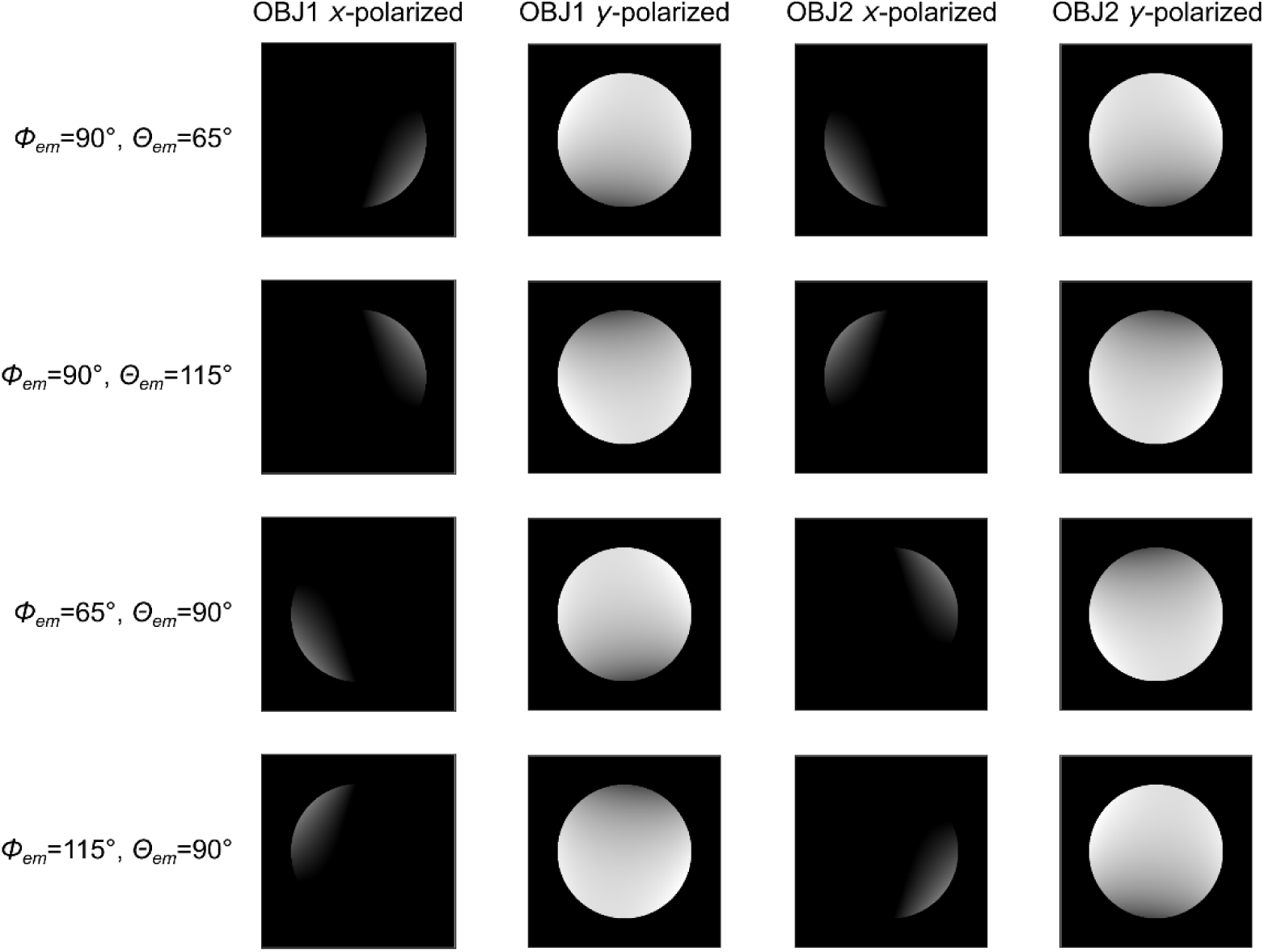
*x*- and *y*-polarized intensity patterns at the back focal planes of two objectives of four degenerate emission dipole orientations (region size 2*λ*_*em*_ × 2*λ*_*em*_, where *λ*_*em*_ is the emission wavelength). The calculation is based on vectorial diffraction theory. See details in Supplemental Note 2. To increase the contrast of images, we assume *NA* = 0.7 for both objectives here.

### Supplemental Note 1. Vectorial diffraction theory calculation of excitation efficiency

The calculation is based on vectorial diffraction theory^1^. Assume the incident laser light has a wavelength of *λ* = 532 nm and is polarized along the *x*-axis; the microscope objective has a numerical aperture *NA* = 0.7 and a focal length *f* = 4 mm. In the dual-view setup the medium filling the space between the objectives and molecules is air. Therefore, it is also reasonable to assume that the laser light is focused into a homogeneous space with refractive index *n* = 1.

Let the incident laser light fill the back-aperture of the objective (filling factor *f*_0_ = 1). The pupil filter is 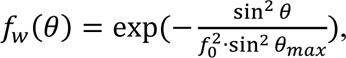 where *θ* is the incident angle and *θ*_*max*_ is the half-aperture angle of the microscope objective defined as 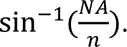 Cylindrical coordinates (*ρ*, *φ*, *z*) are used for focal field calculation for symmetric consideration. The focal field of the Gaussian mode is:

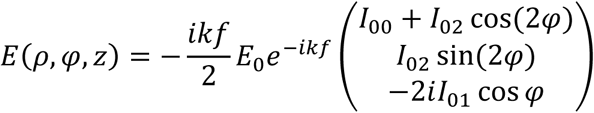

where 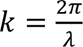 is the wave number, *E*_0_ is a constant electric field amplitude, *I*_00_, *I*_01_, *I*_02_

are integrals:

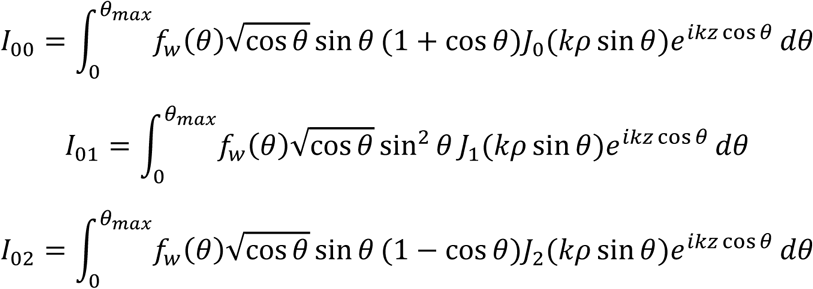

where *J*_*n*_ is the *n*-th order Bessel function of the first kind. The electric field intensity is thus

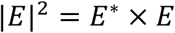

where *E*^∗^ is the complex conjugate of *E*.

The calculated *x*-, *y*- and *z*-polarized electric field intensity distributions at the focal plane are shown in Fig. S3. It is clear that at (or very close to) the focal point the electric field is exactly *x*- polarized, the same as the incident laser light.

Note the calculation assumes *NA* = 0.7. For lower *NA* objectives the focus is looser, so the conclusion that the polarization of electric field at the focal point is the same as that of the linearly polarized excitation light still holds. The excitation efficiency *η*_*abs*_ thus is:

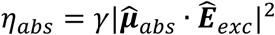

where *γ* is a scalar scaling factor and will be dropped for simplicity.

### Supplemental Note 2. Vectorial diffraction theory calculation of fluorescence collection efficiency

To calculate the fluorescence collection efficiency, we consider the emission dipole as an oscillating electric dipole. In the far-field approximation (*r* ≫ emitting wavelength *λ*), the electric field at a position 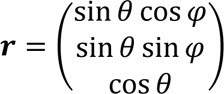 is expressed as:

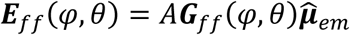

where *A* is a scalar scaling factor and ***G***_*ff*_(*φ*, *θ*) is the Greens’ tensor:

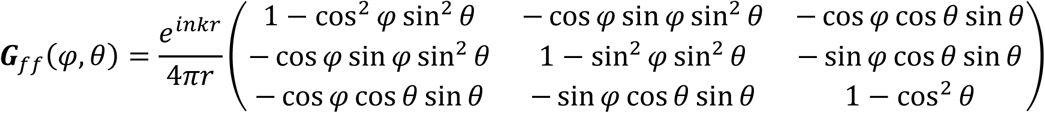

where *n* = 1 is the refractive index of the embedding medium, and 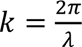 is the wave number.

We assume the objective lens collimates the collected emission light, so that any ray collected will be rotated such that it is parallel to the optical axis of the corresponding objective. In such arrangement, the *s*-polarized component of the electric field remains unchanged, but the *p*- polarized component is rotated so that it is perpendicular to the optical axis. The Green’s tensor at the back focal plane is:

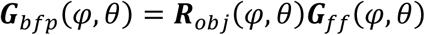

where ***R***_*obj*_(*φ*, *θ*) denotes the act of objective:

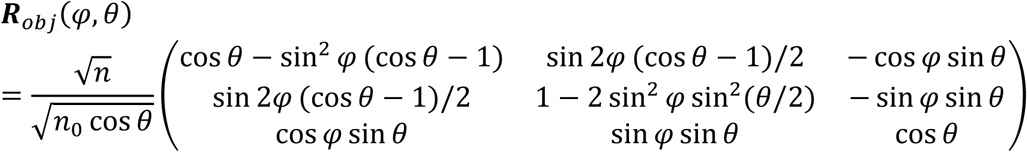

where *n*_0_ = 1 is the refractive index at the back focal plane.

It is more convenient to calculate the electric field at the back focal plane using polar coordinates (*φ*, *ρ*) than spherical coordinates (*φ*, *θ*). The corresponding conversion formulae is:

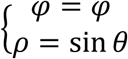

The radial coordinate *ρ* is normalized so that its maximum *ρ*_*max*_ is:

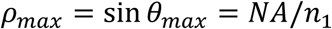

where *θ*_*max*_ is the half-aperture angle of the objective.

The Green’s tensor under polar coordinates becomes:

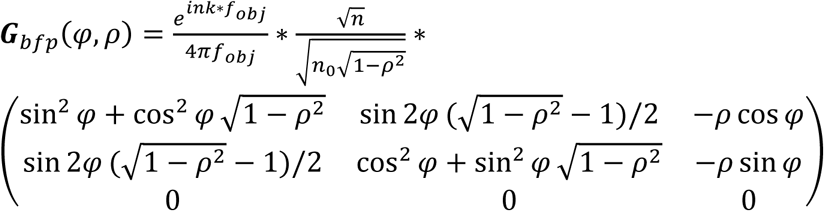

The electric field at the back focal plane is:

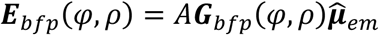

The *x*-polarized component of the electric field at the back focal plane is:

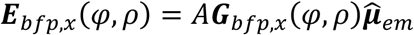

where ***G***_*bfp*,*x*_(*φ*, *ρ*) is the first row of ***G***_*bfp*_(*φ*, *ρ*). The corresponding *x*-polarized intensity is:

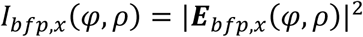

Note that under the rotated coordinate system with respect to each objective, the *s*- and *p*-polarized components of the electric field are exactly *x*- and *y*-polarized components. After passing through the polarizing beam splitter (PBS), the intensity *I*_*x*_ of *x*-polarized component of the electric field recorded by one avalanche photodiode (APD) is calculated by integration:

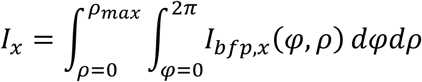

The intensity *I*_*y*_ of *y*-polarized component of the electric field recorded by the other APD is calculated similarly. After integration we find:

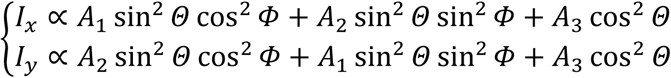

where *A*_1_, *A*_2_, *A*_3_ are functions of *ρ*_*max*_, and therefore, *NA*:

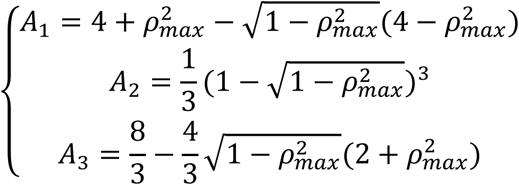

### Supplemental Note 3. Coordinate transformations

The spherical coordinates (*Φ*^′^, *Θ*′) of *μ̂*_*abs*_ under the rotated coordinate system in Fig. S2 can be obtained by applying a transformation matrix corresponding to the right-handed rotation of an angle *Ω* about the *y*-axis to the original coordinates:

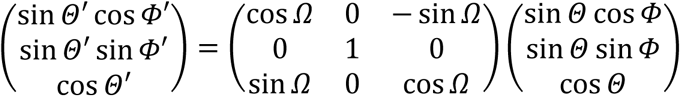

from where we get:

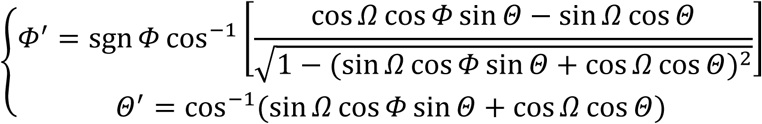

where sgn is the sign function:

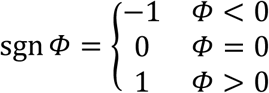

Note the polarization of the linearly polarized light is perpendicular to the optical axis of the objective, so the excitation polarization *Ê*_*exc*_ is fully described by the azimuthal angle *φ*_*exc*_ under the rotated coordinate system. The absorption efficiency *η*_*abs*_ can then be calculated as:

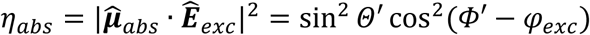

Like the excitation case, we first calculate spherical coordinates (*Φ*^′^, *Θ*′) of the emission dipole orientation *μ̂*_*em*_ under the rotated coordinate system with respect to the objective. After passing through the objective, the intensities of *x*- and *y*-polarized (under the rotated coordinate system) components of emission are found to be:

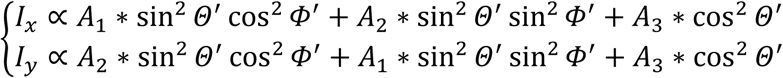

where *A*_1_, *A*_2_, *A*_3_ are functions of *NA* described in Supplemental Note 2.

If avalanche photodiodes (APDs) are placed to detect the *x*- and *y*-polarized emission, the fluorescence collection efficiencies of APDs are proportional to *I*_*x*_ or *I*_*y*_. Therefore, the relative fluorescence collection efficiency of each APD can be fully determined by the knowledge of the emission dipole orientation and *A*_1_, *A*_2_, *A*_3_.

### Supplemental Note 4. Maximum likelihood estimation (MLE) results and Cramér-Rao lower bound (CRLB)

To characterize the spread of dipole orientations from maximum likelihood estimation, we compute the “observed” covariance ellipse^2^ from the covariance matrix and compare it to the expected covariance ellipse computed from the expected Fisher information matrix^3^. Fig. S4 a shows the two ellipses calculated from the example in Fig. 2 a-b in the main text. Fig. S4 b shows the expected and “observed” covariance ellipses of 50 equally distanced orientations on the hemisphere assuming *n*_*tot*_ = 5000, *β* = 4. It is clear that “observed” covariance ellipses almost coincide with the corresponding expected covariance ellipses, indicating MLE results approach the theoretical limit CRLB.

### Supplemental Note 5. 2-fold degeneracy of absorption dipole orientation estimation along the *Φ*_*abs*_ = 0°/180° longitude line

Consider two orientations that (1) their azimuthal angles are the same 0°/180°; (2) their polar angles are complementary under the laboratory coordinate system as shown in Fig. S5. They are in the plane determined by the optical axes of OBJ1 and OBJ2, which is the *xz*-plane under the laboratory coordinate system.

The absorption efficiency *η*_*abs*_ is calculated as:

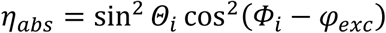

where *Φ*_*i*_, *Θ*_*i*_ (*i* = 1, 2) are the azimuthal and polar angles of the dipole orientation under the rotated coordinate system with respect to OBJ1 or OBJ2.

For the two orientations their *Φ*_*i*_ values are still 0°/180°, so the cos^2^(*Φ*_*i*_ − *φ*_*exc*_) term is the same. Their *Θ*_*i*_ values are either equal or supplementary, but the sin^2^ *Θ*_*i*_ term is still the same. Therefore, *η*_*abs*_ is the same for the two orientations, independent of the *φ*_*exc*_ value, i.e., at each excitation.

### Supplemental Note 6. Excitation polarization modulation in “epi” geometry

To calculate the Fisher information and corresponding CRLB of the excitation polarization modulation method in “epi” geometry, we adapt the formalism developed by Fourkas^4,5^. Since we assume *NA* = 0.5 for both objectives in the dual-view setup, the effective *NA* of the system is ∼0.68. To make a fair comparison, we assume an air objective of *NA* = 0.7 is used in the excitation polarization modulation method in “epi” geometry. We excite the molecule at the focus 4 times with *φ*_*exc*_ = 0°, 45°, 90°, 135° respectively. At each excitation the absorption efficiency is calculated as:

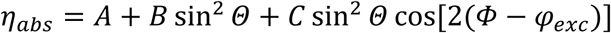

where the *A*, *B*, *C* terms are:

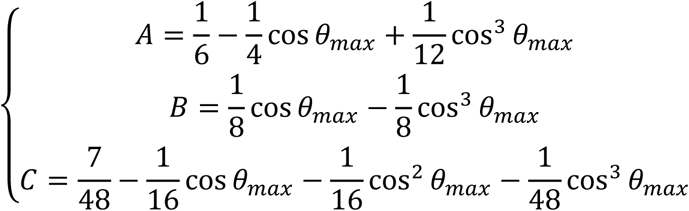

where 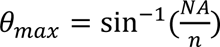 (refractive index *n* = 1).

We calculate CRLBs of azimuthal and polar angles in single-molecule absorption dipole orientation estimation for excitation polarization modulation method in “epi” and dual-view geometry. See details in Methods. The calculation is performed on 10,000 equally spaced orientations on the unit sphere assuming *n*_*tot*_ = 5000. The results at signal-to-background ratio *β* = ∞ (i.e., background-free case) are shown in Fig. 2 h-i in the main text. The results at signal-to-background ratio *β* = 4 are shown in Fig. S6.

### Supplemental Note 7. 4-fold degeneracy of emission dipole orientation estimation

The 4-fold degeneracy of emission dipole orientation estimation comes from the complementary geometry in our setup. Given an emission dipole orientation ground truth, one degenerate orientation is symmetric to the ground truth about the *y*-axis of laboratory coordinate system; the other two degenerate orientations are symmetric to the ground truth about the planes perpendicular to the optical axes of two objectives. See an example in Fig. S7. Given one emission dipole orientation, its degenerate orientations can be readily calculated by coordinate transformations.

### Supplemental Note 8. Characterizing emission dipole orientation estimation results

For each emission dipole orientation ground truth, the simulation and estimation process are repeated multiple times. Every time the emission dipole orientation is estimated, the other three degenerate orientations can be calculated by geometry (Supplemental Note 7). The estimated emission dipole orientations usually form 4 clusters. In some regions of the parameter space cluster merging occurs, resulting in 2 or 1 clusters.

We calculate the angle *δ*_*i*_ between each estimated emission dipole orientation *μ̂*_*i*_ and emission dipole orientation ground truth *μ̂*_*em*_:

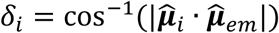

To characterize the emission dipole orientation estimation, we select the estimated emission dipole orientations if *δ*_*i*_ ≤ *δ*_*thershold*_, where *δ*_*thershold*_ is a threshold value. We set *δ*_*thershold*_ = 10° here.

In another scenario, after picking one of the four candidates as estimated emission dipole orientation, we define the “hit” rate as the ratio of the number of estimated emission dipole orientations whose angle with the corresponding emission dipole ground truth is smaller than *δ*_*thershold*_ to the total number of estimated emission dipole orientations.

## REFERENCES

(1) Forkey, J. N.; Quinlan, M. E.; Goldman, Y. E. Protein Structural Dynamics by Single-Molecule Fluorescence Polarization. Prog. Biophys. Mol. Biol. 2000, 74 (1), 1–35. 10.1016/S0079-6107(00)00015-8.

(2) Backlund, M. P.; Lew, M. D.; Backer, A. S.; Sahl, S. J.; Moerner, W. E. The Role of Molecular Dipole Orientation in Single-Molecule Fluorescence Microscopy and Implications for Super-Resolution Imaging. ChemPhysChem 2014, 15 (4), 587–599. 10.1002/cphc.201300880.

(3) Brasselet, S.; Alonso, M. A. Polarization Microscopy: From Ensemble Structural Imaging to Single-Molecule 3D Orientation and Localization Microscopy. Optica 2023, 10 (11), 1486–1510. 10.1364/OPTICA.502119.

(4) Zhang, O.; Guo, Z.; He, Y.; Wu, T.; Vahey, M. D.; Lew, M. D. Six-Dimensional Single-Molecule Imaging with Isotropic Resolution Using a Multi-View Reflector Microscope. Nat. Photonics 2022. 10.1038/s41566-022-01116-6.

(5) Bruggeman, E.; Zhang, O.; Needham, L.-M.; Körbel, M.; Daly, S.; Cheetham, M.; Peters, R.; Wu, T.; Klymchenko, A. S.; Davis, S. J.; Paluch, E. K.; Klenerman, D.; Lew, M. D.; O’Holleran, K.; Lee, S. F. POLCAM: Instant Molecular Orientation Microscopy for the Life Sciences. Nat. Methods 2024, 1–11. 10.1038/s41592-024-02382-8.

(6) Zhang, O.; Lew, M. D. Single-Molecule Orientation-Localization Microscopy: Applications and Approaches. Q. Rev. Biophys. 2024, 57, e17. 10.1017/S0033583524000167.

(7) Lakowicz, J. R. Principles of Fluorescence Spectroscopy; Springer Science: New York, 2006.

(8) Sick, B.; Hecht, B.; Novotny, L. Orientational Imaging of Single Molecules by Annular Illumination. Phys. Rev. Lett. 2000, 85 (21), 4482–4485. 10.1103/PhysRevLett.85.4482.

(9) Prummer, M.; Sick, B.; Hecht, B.; Wild, U. P. Three-Dimensional Optical Polarization Tomography of Single Molecules. J. Chem. Phys. 2003, 118 (21), 9824–9829. 10.1063/1.1569848.

(10) Bartko, A. P.; Dickson, R. M. Imaging Three-Dimensional Single Molecule Orientations. J. Phys. Chem. B 1999, 103 (51), 11237–11241. 10.1021/jp993364q.

(11) Hohlbein, J.; Hübner, C. G. Simple Scheme for Rapid Three-Dimensional Orientation Determination of the Emission Dipole of Single Molecules. Appl. Phys. Lett. 2005, 86 (12), 121104. 10.1063/1.1888040.

(12) Böhmer, M.; Enderlein, J. Orientation Imaging of Single Molecules by Wide-Field Epifluorescence Microscopy. JOSA B 2003, 20 (3), 554–559. 10.1364/JOSAB.20.000554.

(13) Sikorski, Z.; Davis, L. M. Engineering the Collected Field for Single-Molecule Orientation Determination. Opt. Express 2008, 16 (6), 3660–3673. 10.1364/OE.16.003660.

(14) Stelzer, E. H. K.; Lindek, S. Fundamental Reduction of the Observation Volume in Far-Field Light Microscopy by Detection Orthogonal to the Illumination Axis: Confocal Theta Microscopy. Opt. Commun. 1994, 111 (5), 536–547. 10.1016/0030-4018(94)90533-9.

(15) Wu, Y.; Ghitani, A.; Christensen, R.; Santella, A.; Du, Z.; Rondeau, G.; Bao, Z.; Colón-Ramos, D.; Shroff, H. Inverted Selective Plane Illumination Microscopy (iSPIM) Enables Coupled Cell Identity Lineaging and Neurodevelopmental Imaging in Caenorhabditis Elegans. Proc. Natl. Acad. Sci. U. S. A. 2011, 108 (43), 17708–17713. 10.1073/pnas.1108494108.

(16) Wu, Y.; Wawrzusin, P.; Senseney, J.; Fischer, R. S.; Christensen, R.; Santella, A.; York, A. G.; Winter, P. W.; Waterman, C. M.; Bao, Z.; Colón-Ramos, D. A.; Mcauliffe, M.; Shroff, H. Spatially Isotropic Four-Dimensional Imaging with Dual-View Plane Illumination Microscopy. Nat. Biotechnol. 2013, 31 (11), 1032–1038. 10.1038/nbt.2713.

(17) Wu, Y.; Chandris, P.; Winter, P. W.; Kim, E. Y.; Jaumouillé, V.; Kumar, A.; Guo, M.; Leung, J. M.; Smith, C.; Rey-Suarez, I.; Liu, H.; Waterman, C. M.; Ramamurthi, K. S.; Riviere, P. J. L.; Shroff, H. Simultaneous Multiview Capture and Fusion Improves Spatial Resolution in Wide-Field and Light-Sheet Microscopy. Optica 2016, 3 (8), 897–910. 10.1364/OPTICA.3.000897.

(18) Stelzer, E. H. K.; Strobl, F.; Chang, B.-J.; Preusser, F.; Preibisch, S.; McDole, K.; Fiolka, R. Light Sheet Fluorescence Microscopy. Nat. Rev. Methods Primer 2021, 1 (1), 1–25. 10.1038/s43586-021-00069-4.

(19) Wu, Y.; Han, X.; Su, Y.; Glidewell, M.; Daniels, J. S.; Liu, J.; Sengupta, T.; Rey-Suarez, I.; Fischer, R.; Patel, A.; Combs, C.; Sun, J.; Wu, X.; Christensen, R.; Smith, C.; Bao, L.; Sun, Y.; Duncan, L. H.; Chen, J.; Pommier, Y.; Shi, Y.-B.; Murphy, E.; Roy, S.; Upadhyaya, A.; Colón-Ramos, D.; La Riviere, P.; Shroff, H. Multiview Confocal Super-Resolution Microscopy. Nature 2021, 600 (7888), 279–284. 10.1038/s41586-021-04110-0.

(20) Chandler, T.; Mehta, S.; Shroff, H.; Oldenbourg, R.; Rivière, P. J. L. Single-Fluorophore Orientation Determination with Multiview Polarized Illumination: Modeling and Microscope Design. Opt. Express 2017, 25 (25), 31309–31325. 10.1364/OE.25.031309.

(21) Thorsen, R. Ø.; Hulleman, C. N.; Rieger, B.; Stallinga, S. Photon Efficient Orientation Estimation Using Polarization Modulation in Single-Molecule Localization Microscopy. Biomed. Opt. Express 2022, 13 (5), 2835. 10.1364/boe.452159.

(22) Chandler, T.; Guo, M.; Su, Y.; Chen, J.; Wu, Y.; Liu, J.; Agashe, A.; Fischer, R. S.; Mehta, S. B.; Kumar, A.; Baskin, T. I.; Jaumouillé, V.; Liu, H.; Swaminathan, V.; Nain, A. S.; Oldenbourg, R.; La Riviere, P. J.; Shroff, H. Volumetric Imaging of the 3D Orientation of Cellular Structures with a Polarized Fluorescence Light-Sheet Microscope. Proc. Natl. Acad. Sci. 2025, 122 (8), e2406679122. 10.1073/pnas.2406679122.

(23) Backer, A. S.; Moerner, W. E. Extending Single-Molecule Microscopy Using Optical Fourier Processing. J. Phys. Chem. B 2014, 118 (28), 8313–8329. 10.1021/jp501778z.

(24) Zhang, O.; Zhou, W.; Lu, J.; Wu, T.; Lew, M. D. Resolving the Three-Dimensional Rotational and Translational Dynamics of Single Molecules Using Radially and Azimuthally Polarized Fluorescence. Nano Lett. 2022, 22 (3), 1024–1031. 10.1021/acs.nanolett.1c03948.

(25) Levitus, M.; Negri, R. M.; Aramendia, P. F. Rotational Relaxation of Carbocyanines. Comparative Study with the Isomerization Dynamics. J. Phys. Chem. 1995, 99 (39), 14231–14239. 10.1021/j100039a008.

(26) Synak, A.; Bojarski, P. Transition-Moment Directions of Selected Carbocyanines from Emission Anisotropy and Linear Dichroism Measurements in Uniaxially Stretched Polymer Films. Chem. Phys. Lett. 2005, 416 (4), 300–304. 10.1016/j.cplett.2005.09.097.

(27) Sanborn, M. E.; Connolly, B. K.; Gurunathan, K.; Levitus, M. Fluorescence Properties and Photophysics of the Sulfoindocyanine Cy3 Linked Covalently to DNA. J. Phys. Chem. B 2007, 111 (37), 11064–11074. 10.1021/jp072912u.

(28) Fourkas, J. T. Rapid Determination of the Three-Dimensional Orientation of Single Molecules. Opt Lett 2001, 26 (4), 211–213. 10.1364/OL.26.000211.

(29) Beckwith, J. S.; Yang, H. Information Bounds in Determining the 3D Orientation of a Single Emitter or Scatterer Using Point-Detector-Based Division-of-Amplitude Polarimetry. J. Chem. Phys. 2021, 155 (14), 144110. 10.1063/5.0065034.

(30) Kawski, A.; Gryczyński, Z. Determination of the Transition-Moment Directions from Photoselection in Partially Oriented Systems. Z. Für Naturforschung A 1987, 42 (8), 808–812. 10.1515/zna-1987-0807.

(31) Ha, T.; Laurence, T. a; Chemla, D. S.; Weiss, S. Polarization Spectroscopy of Single Fluorescent Molecules. J Phys Chem B 1999, 103 (33), 6839–6850. 10.1021/jp990948j.

(32) Luschtinetz, F.; Dosche, C.; Kumke, M. U. Influence of Streptavidin on the Absorption and Fluorescence Properties of Cyanine Dyes. Bioconjug. Chem. 2009, 20 (3), 576–582. 10.1021/bc800497v.

(33) Mukerjee, A.; Sørensen, T. J.; Ranjan, A. P.; Raut, S.; Gryczynski, I.; Vishwanatha, J. K.; Gryczynski, Z. Spectroscopic Properties of Curcumin: Orientation of Transition Moments. J. Phys. Chem. B 2010, 114 (39), 12679–12684. 10.1021/jp104075f.

(34) Curcio, V.; Alemán-Castañeda, L. A.; Brown, T. G.; Brasselet, S.; Alonso, M. A. Birefringent Fourier Filtering for Single Molecule Coordinate and Height Super-Resolution Imaging with Dithering and Orientation. Nat. Commun. 2020, 11 (1), 5307. 10.1038/s41467-020-19064-6.

(35) Hulleman, C. N.; Thorsen, R. Ø.; Kim, E.; Dekker, C.; Stallinga, S.; Rieger, B. Simultaneous Orientation and 3D Localization Microscopy with a Vortex Point Spread Function. Nat. Commun. 2021, 12 (1), 5934. 10.1038/s41467-021-26228-5.

(36) Chizhik, A. I.; Chizhik, A. M.; Huss, A.; Jäger, R.; Meixner, A. J. Nanoscale Probing of Dielectric Interfaces with Single-Molecule Excitation Patterns and Radially Polarized Illumination. J. Phys. Chem. Lett. 2011, 2 (17), 2152–2157. 10.1021/jz200934y.

(37) Karedla, N.; Stein, S. C.; Hähnel, D.; Gregor, I.; Chizhik, A.; Enderlein, J. Simultaneous Measurement of the Three-Dimensional Orientation of Excitation and Emission Dipoles. Phys. Rev. Lett. 2015, 115 (17), 173002. 10.1103/PhysRevLett.115.173002.

(38) Backlund, M. P.; Lew, M. D.; Backer, A. S.; Sahl, S. J.; Grover, G.; Agrawal, A.; Piestun, R.; Moerner, W. E. Simultaneous, Accurate Measurement of the 3D Position and Orientation of Single Molecules. Proc. Natl. Acad. Sci. 2012, 109 (47), 19087–19092. 10.1073/pnas.1216687109.

(39) Dahlberg, P. D.; Moerner, W. E. Cryogenic Super-Resolution Fluorescence and Electron Microscopy Correlated at the Nanoscale. Annu Rev Phys Chem 2021, 72 (1), 253–278. 10.1146/annurev-physchem-090319-051546.

## References

(1) Novotny, L.; Hecht, B. Principles of Nano-Optics, 2nd ed.; Cambridge University Press: Cambridge, 2012. 10.1017/CBO9780511794193.

(2) Balzarotti, F.; Eilers, Y.; Gwosch, K. C.; Gynnå, A. H.; Westphal, V.; Stefani, F. D.; Elf, J.; Hell, S. W. Nanometer Resolution Imaging and Tracking of Fluorescent Molecules with Minimal Photon Fluxes. Science 2017, 355 (6325), 606–612. 10.1126/science.aak9913.

(3) Coe, D. Fisher Matrices and Confidence Ellipses: A Quick-Start Guide and Software. arXiv June 23, 2009. 10.48550/arXiv.0906.4123.

(4) Fourkas, J. T. Rapid Determination of the Three-Dimensional Orientation of Single Molecules. Opt Lett 2001, 26 (4), 211–213. 10.1364/OL.26.000211.

(5) Beckwith, J. S.; Yang, H. Information Bounds in Determining the 3D Orientation of a Single Emitter or Scatterer Using Point-Detector-Based Division-of-Amplitude Polarimetry. J. Chem. Phys. 2021, 155 (14), 144110. 10.1063/5.0065034.

